# The Arp2/3 complex is required for *in situ* haptotactic response of microglia to iC3b

**DOI:** 10.1101/2025.05.21.655384

**Authors:** Summer G. Paulson, Isabella Swafford, Fritz W. Lischka, Jeremy D. Rotty

**Affiliations:** Uniformed Services University of the Health Sciences, Department of Biochemistry, Bethesda, MD, USA, 20814; Uniformed Services University of the Health Sciences, Department of Anatomy, Physiology, and Genetics, Bethesda, MD, USA, 20814; The Henry M. Jackson Foundation for the advancement of Military Medicine, Bethesda, MD, USA, 20817; Uniformed Services University of the Health Sciences, Biomedical Instrumentation Center, Bethesda, MD, USA, 20814

**Keywords:** Microglia, iC3b, Arp2/3 complex, haptotaxis, synaptic pruning, confined motility, phagocytosis

## Abstract

Microglia maintain brain homeostasis by performing iC3b-mediated synaptic pruning on excessive dendritic spines during neurodevelopment. Cellular interaction with iC3b is mediated by the Arp2/3 complex in other cell types, but understudied in microglia. Using a combination of *in vitro* and *in situ* physical confinement studies, we examined CR3-dependent clearance of iC3b in microglia and the role the Arp2/3 complex plays in enabling this clearance. We demonstrated Arp2/3 complex inhibition decreased phagocytosis and cell motility *in vitro*. Furthermore, we demonstrate that microglia-like cells are able to remove immobilized iC3b from the substrate in an Arp2/3-dependent fashion, in a process reminiscent of trogocytic synaptic pruning. We also used a novel approach to immobilize an iC3b gradient onto a substrate and demonstrate Arp2/3-dependent haptotactic migration toward increasing iC3b concentrations. Microglia demonstrate a persistent inability to stably interact with iC3b-coated beads in hippocampal slice cultures upon Arp2/3 complex inhibition. As a whole, the present study establishes new approaches to systematically interrogate molecular pathways relevant to synaptic pruning, advances the understanding of iC3b phagocytosis as a haptotactic response, and confirms that the Arp2/3-dependent haptotactic response is relevant for microglia in their normal physiological microenvironment.

## INTRODUCTION

Microglia are the resident immune cells of the central nervous system (CNS) (*1*, *2*). They respond to a variety of phagocytic markers, including the complement proteins iC3b and C1q, that bind to and label phagocytic targets upon activation of the complement cascade (*3*). These factors label and clear foreign bodies and debris but also have been implicated in dendritic spine pruning in the CNS (*4–6*). Dendritic spines, located along the length of neuronal dendrites, are the postsynaptic sites of synapses, a docking point to facilitate signal relay between neurons (*7*). During development, neurons initially overpopulate their dendrites with new dendritic spines (*8*). Excessive spines are then marked for microglial phagocytosis using iC3b (*9*). Microglia respond to the iC3b cue via complement receptor 3 (CR3, also known as Mac1) (*10*). This results in synaptic pruning and remodeling of the neuronal network by microglia (*11*).

The CR3 receptor is composed of 2 integrins: CD11b (αM integrin) and CD18 (β2 integrin) (*12*). The αMβ2 heterodimer binds directly to phagocytic cues such as iC3b and initiates the process of internalization via mechanosensitive clutch formation and Arp2/3 complex-dependent membrane protrusion (*13*). Integrins are also vital in adhesion-dependent mesenchymal cell migration by similarly linking the surrounding extracellular matrix (ECM) to adhesive structures and the actomyosin machinery in cells (*14*). The seven-subunit Arp2/3 complex generates dense branched actin networks in response to integrin engagement in the context of migration and phagocytosis (*15*). Upon docking to a pre-existing ‘mother’ actin filament, Arp2/3 complex polymerizes a branched ‘daughter’ filament at a characteristic 70° angle (*16*). The growing barbed (+) ends of these branched actin arrays provide the physical force that pushes against the membrane, causing characteristic dynamic protrusions at the cell edge called lamellipodia (*16*).

Arp2/3 complex-deficient cells fail to generate iC3b-responsive phagocytic cups and cannot sense fibronectin gradients, suggesting a fundamental defect related to integrin sensing (*17*). However, it is not yet known whether the Arp2/3 complex is necessary for complement-mediated phagocytosis *in vivo*, such as in the CNS during synaptic pruning.

Complement-mediated phagocytosis has been studied extensively in macrophages, examining phagocytic cup formation, macrophage response to complement signaling, and downstream cytokine production (*18*, *19*). However, much of this work has been done in immortalized macrophage lines *in vitro* in 2D culture experiments that do not consider how tissue confinement or interaction with the complex set of cues in the endogenous microenvironment might modulate this process. Furthermore, insights generated from studying macrophage complement-mediated phagocytosis may not directly translate to microglia as these cell types are vastly different, despite microglia’s common definition as the macrophages of the central nervous system. A study examining gene expression differences between microglia and monocyte-derived macrophages found 239 genes specifically expressed in microglia (*20*). In addition, microglia have a dramatically different morphology and *in vivo* behavior compared to tissue macrophages (*20*). Prior research on the Arp2/3 complex within the CNS has focused on how neuronal Arp2/3 complex supports dendritic spine generation and morphology (*21*). The role of the Arp2/3 complex within microglia, and how it responds to specific environmental cues remains understudied in comparison.

In the present work, we use a combination of *in vitro* and *in situ* physical confinement studies to interrogate CR3-dependent clearance of iC3b in microglia. Arp2/3 complex inhibition decreased phagocytosis and cell motility *in vitro*. Furthermore, we demonstrate that microglia-like cells are able to remove immobilized iC3b from the substrate in an Arp2/3-dependent fashion, in a process reminiscent of trogocytic synaptic pruning. Trogocytosis is the act of one cell phagocytizing only a small portion of another cells, differing from phagocytosis due to the way engulfment occurs, but still implemented with need for the Arp2/3 complex (*22*, *23*). We also used a novel approach to immobilize an iC3b gradient onto a substrate and demonstrate Arp2/3-dependent haptotactic migration toward increasing iC3b concentrations. Finally, we quantified microglial response to ATP and iC3b-labeled beads in murine hippocampal slice culture to interrogate extracellular sensing within the physiological microenvironment. In line with a significant body of *in vitro* work, we demonstrate that Arp2/3 complex inhibition does not impact the microglial response to the chemotactic ligand ATP. However, microglia demonstrate a persistent inability to stably interact with iC3b-coated beads upon Arp2/3 complex inhibition. As a whole, the present study establishes new approaches to systematically interrogate molecular pathways relevant to synaptic pruning, advances the understanding of iC3b phagocytosis as a haptotactic response, and confirms that the Arp2/3-dependent haptotactic response is relevant for microglia in their normal physiological microenvironment.

## RESULTS

### iC3b acts as a phagocytic cue independent of how cells interact with it

iC3b, a phagocytic “eat me” cue common to the brain and CNS, represents a microenvironmental cue that stimulates a specific microglial response in the context of synaptic pruning. To facilitate our assessment of microglia-iC3b interactions, we labeled purified human iC3b with Alexa Fluor 555 (denotated as AF-iC3b). During pilot studies using the iC3b to coat dishes, we noticed depositions of fluorescent puncta in the AF-iC3b wells. We confirmed that these puncta were wholly composed of iC3b and not aggregated fluorescent dye by demonstrating that fluorescent signal correlated tightly with purified iC3b (**Supplemental Figure S1A**). To confirm the specificity of the AF-iC3b interaction with CR3 in our cells, we performed adhesion assay after inhibiting the CR3 receptor using blocking antibodies against both integrin partners. BV-2 cells (a microglia-like cell line) were incubated with FBS (negative control), IgG (a negative isotype control), αM-, or β2-blocking antibodies before seeding onto either 10µg/mL fibronectin (FN) or 20µg/mL AF-iC3b coated wells. After waiting 2 hours for cells to adhere, each well was washed and imaged (**Supplemental Figure S1B**). A significant reduction in adhesion to AF-iC3b occurred when either αM or β2 integrin was inhibited, while adhesion to FN remained unchanged by CR3 inhibition (**Supplemental Figure S1C, S1D**). Alternatively, there was no significant difference in the total number of cells present before the wash when comparing within antibody plating groups (**Supplemental Figure S1E**). These findings gave us confidence that AF-iC3b effectively coats glass and plastic, and that its interaction with our cells is CR3-dependent.

Previous research in the lab examined how different ECM coatings affect cell migration (*24*). We hypothesized that iC3b similarly induces a migratory response. BV-2 cells were plated on 10µg/mL FN or on 1µg/mL, 10µg/mL, or 20µg/mL of AF-iC3b. Compared to FN, cells migrated much more slowly on AF-iC3b (**Figure 1A**), with a dose-dependent decrease in accumulated distance traveled across the AF-iC3b concentrations and compared to FN (**Figure 1B**). Lastly, cells plated on the AF-iC3b were less persistent, following the overall trend demonstrated by velocity measurements (**Figure 1C**). Together these results establish that microglia adhere more strongly to AF-iC3b than to FN. Therefore, we decided to examine whether these results were due to cells removing AF-iC3b from the surface of the well rather than using it primarily as a migration substrate.

**Figure 1:**
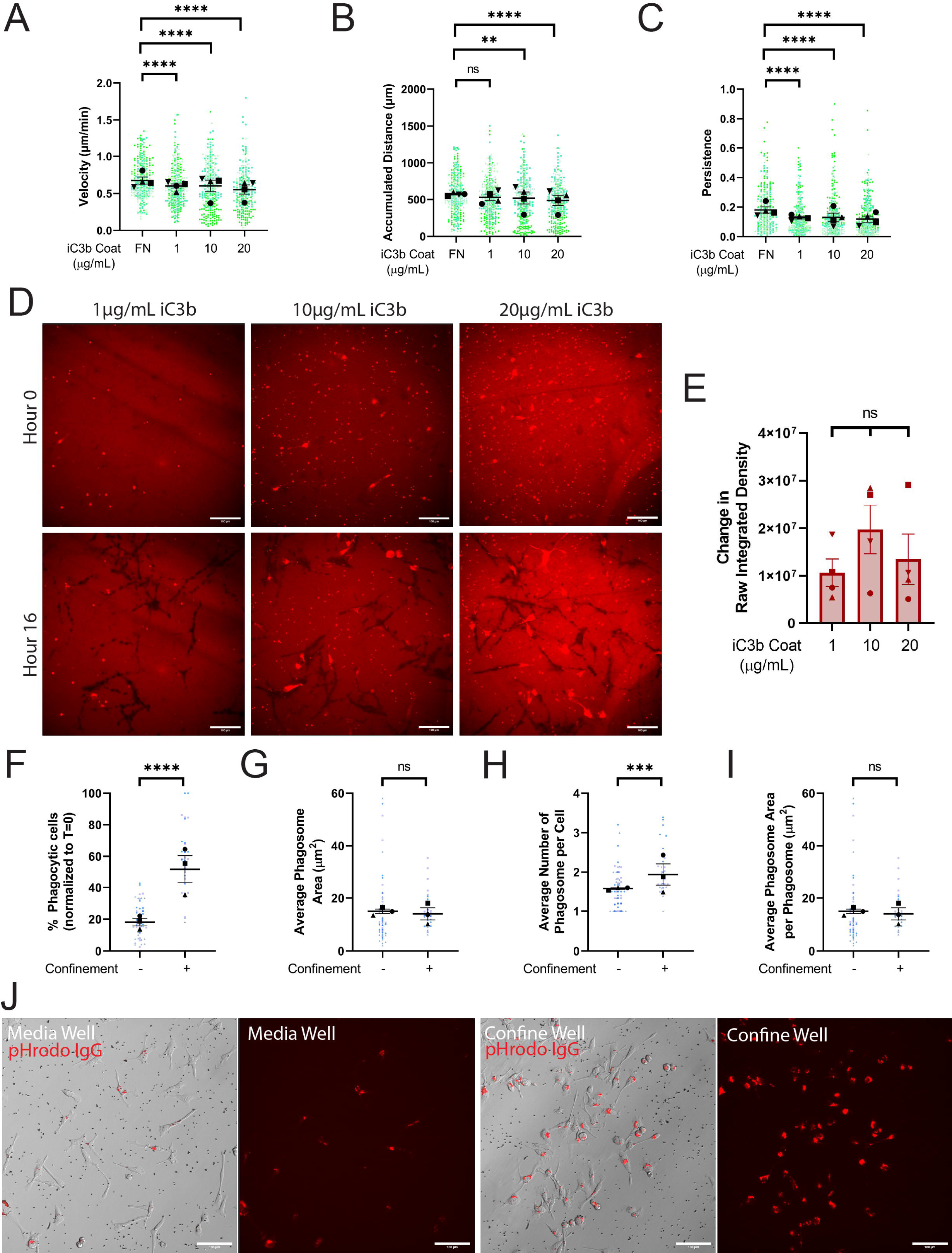
iC3b phagocytosis enhanced by confinement, not iC3b dosage. A-E) Cells plated on either 10µg/mL FN, 1µg/mL AF-iC3b, 10µg/mL AF-iC3b, or 20µg/mL AF-iC3b. A) Velocity of cells migrating (µm/min). B) The maximum accumulated distance (µm) that the cell travelled. C) Persistence of the cell during the length of its track. D) Example images at each AF-iC3b coating dosage demonstrating phagocytosis of coating after 16 hours. Scale bar represents 100µm. E) Quantification of amount of iC3b phagocytized off bottom of the dish, demonstrated in change in raw integrated density of fluorescence. F-J) Cells plated on either in cell culture media or confined under 1% agarose and introduced to pHrodo-iC3b-opsonized beads. F) The percentage of fluorescent cells in a field of view, comparing unconfined and confined cells, normalized to T = 0 hours. Any cell fluorescent at T = 0 was removed from further counting. G) The average phagosome area per phagocytic cell (µm^2^). This is calculated by dividing the average size of all phagosomes in a field of view by the number of fluorescent cells counted (denoted phagocytic cells). H) Average number of phagosomes per phagocytic cell. I) Average phagosome size (µm^2^). This is the average size of phagosomes per field of view divided by the average number of phagosomes per field of view. J) Example composite images of phase contrast and pHrodo-red merged images 2 hours after pHrodo-iC3b beads were added to media wells (left) or confined agarose wells (right). The pHrodo-red signal from these merges has been isolated alongside the merge to emphasize internalized pHrodo-iC3b-bead signal. Scale bar represents 100µm. For all graphs, black points demonstrate experiment means, colored points demonstrate individual cell values for each run. N = 4 experiments for migration (A-E) or N = 3 experiments for phagocytosis (F-J); n = 50 cells for each condition per experiment for migration (A-C), n = 10 fields of view per condition for coating phagocytosis (E), or n = 15 fields of view for each condition per experiment for phagocytosis (F-J). For (A-E), statistical analysis was assessed using via Kruskal–Wallis test with Dunn multiple comparisons: ns = not significant, **p < 0.01, ****p < 0.0001. For (F-J), statistical analysis was assessed using the Mann–Whitney tests: ns = not significant, **p < 0.01, ***p < 0.001, ****p < 0.0001. Error Bars represent SEM.

Creating binary masks and measuring the raw integrated density of the entire field of view, we examined differences between the beginning and end of the 16-hour run to determine how much AF-iC3b had been removed. We observed a large amount of the coating was being pulled up and phagocytized, regardless of the surface concentration of AF-iC3b (**Figure 1D, quantified in Figure 1E**). As a result, cells became fluorescent as they ingested the AF-iC3b, demonstrating that they were internalizing the label. It is reasonable to conclude that the decreased migratory values on AF-iC3b compared to FN are due to persistent cell interaction with substrate-associated AF-iC3b. Next, we decided to explicitly investigate phagocytosis with pHrodo-iC3b-opsonized 2-micron polyspheres (here on referred to as beads).

The 2-micron bead size was chosen due to its similarity to the size of iC3b-tagged dendritic spines that microglia would interact with during synaptic pruning. The pHrodo tag fluoresces under acidic conditions, such as within the phagolysosome, allowing for unbiased detection of complete phagocytosis (**Supplemental Figure S2A**). To rule out any signal from external beads, we compared external and internalized beads and demonstrated a clear difference between these two states (**Supplemental Figure S2B**). While modeling iC3b-dependent synaptic pruning in an easily controlled *in vitro* environment, we decided to also incorporate other CNS microenvironmental cues to determine whether they affected phagocytosis. We utilized an under-agarose confinement system (*25*), which was originally inspired by Heit and Kubes (*26*) (**Supplemental Figure S2A**). With this assay, cells were either in wells filled with cell culture media (Media Well) or confined underneath a layer of 1% agarose (Confined Well). Because of this use of confinement, normalizing the number of beads present became another essential control point to rule out population differences due simply to altered local bead availability (**Supplemental Figure S2C-D**). This is a factor primarily in interpreting under agarose data, as beads injected under the agarose do not disperse evenly like they do in media. Due to this variability between confined fields of view, we calculated the number of beads in each field of view and categorized them into different density groups. As the majority of media well fields of view were categorized as “low density” (**Supplemental Figure S2D, bottom**), confined fields of view in the same category (**Supplemental Figure S2D, middle**) were exclusively used for analysis.

Phagocytosis of pHrodo-iC3b-beads was measured at 2 hours, before the cells became saturated with beads. Phagocytosis increased significantly under confinement, with a greater than 2-fold increase in the percentage of phagocytic cells compared to media wells (**Figure 2F**). This increase in phagocytic cells also reflected an increase in overall bead consumption per cell, as indicated by the increased average phagosome area and number of phagosomes per phagocytic cell under confinement (**Figure 1G-H**). Surprisingly, we observed no increase in the average size of phagosomes, taken by dividing the phagosome area in the field of view by the number of phagosomes in the field of view (**Figure 1I**). Thus, cells traffic beads to new phagosomes rather than building fewer, larger phagosomes. This is also visible in **Figure 1J**, where comparison of just the pHrodo images demonstrates similarly sized phagosomes for the media and confined wells. These data confirm that BV2 microglia-like cells interact with iC3b in two different contexts (on the surface of a dish, and opsonized onto beads), and in unconfined and confined settings. Next, we wanted to interrogate whether the Arp2/3 complex was required for these responses.

**Figure 2:**
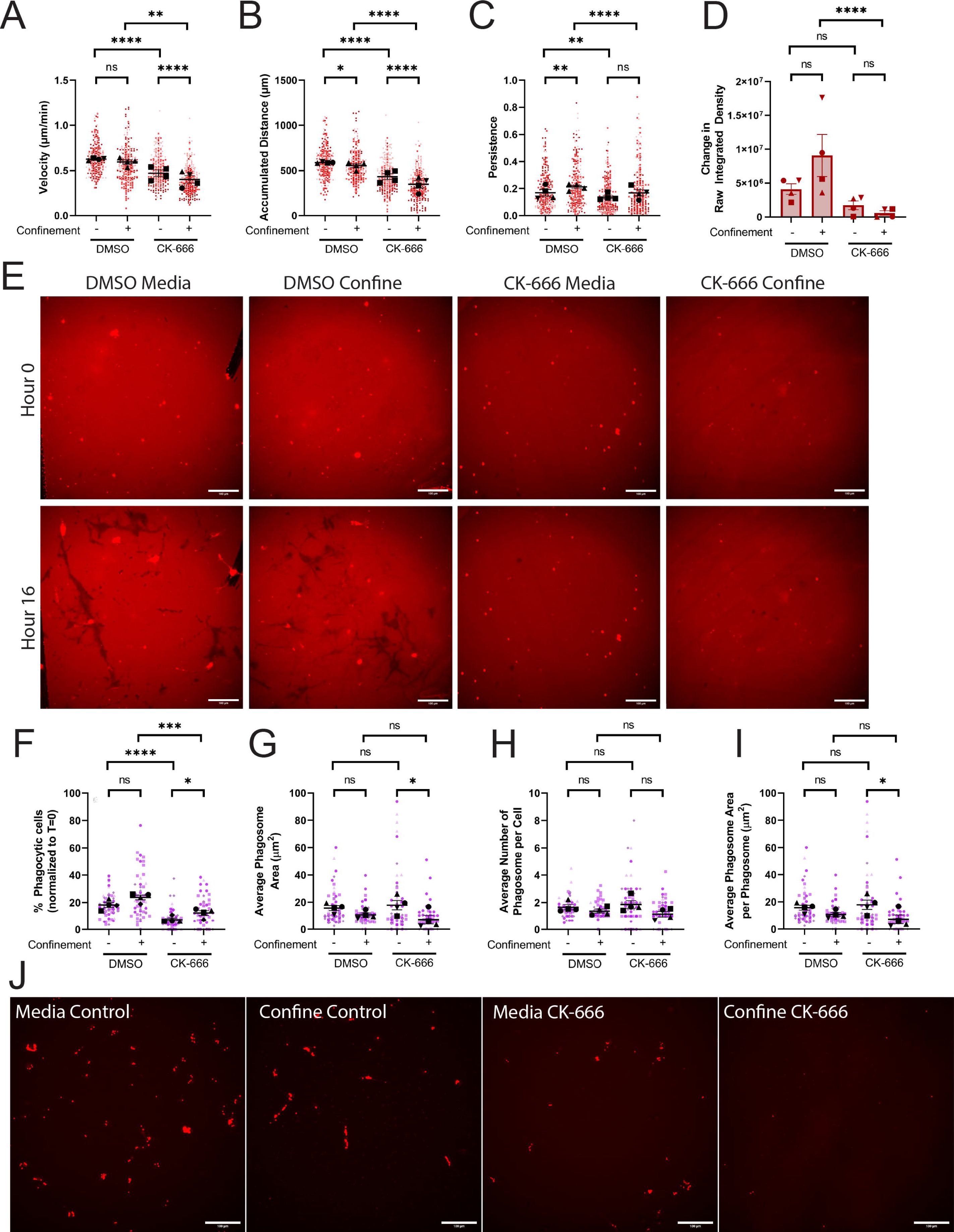
The Arp2/3 complex is required for iC3b phagocytosis, but confinement rescue is context dependent. A-E) Cells plated on 10µg/mL AF-iC3b in each condition: Media control (vehicle – DMSO), Confined control (1% Agarose) plus Vehicle, Media control plus CK-666, Confined (1% Agarose) plus CK-666. A) Velocity of cells migrating (µm/min). B) The maximum accumulated distance (µm) that the cell travelled. C) Persistence of the cell during the length of its track. D) Quantification of amount of iC3b phagocytized off bottom of the dish, demonstrated in change in raw integrated density of fluorescence. E) Example images of phagocytosis of AF-iC3b coating under each treatment condition after 16 hours. Scale bar represents 100µm. F-J) Cells plated on 10µg/mL FN in each condition and given pHrodo-iC3b beads: Media control (vehicle), Confined control (1% Agarose) plus Vehicle, Media control plus CK-666, Confined (1% Agarose) plus CK-666. F) The percentage of fluorescent cells in a field of view, comparing unconfined and confined cells, normalized to T = 0 hours. Any cell fluorescent at T = 0 was removed from further counting. G) The average phagosome area per phagocytic cell (µm^2^). This is calculated by dividing the average size of all phagosomes in a field of view by the number of fluorescent cells counted (denoted phagocytic cells). H) Average number of phagosomes per phagocytic cell. I) Average phagosome size (µm^2^). This is the average size of phagosomes per field of view divided by the average number of phagosomes per field of view. J) Example composite images of phase contrast and pHrodo-red merged images 2 hours after pHrodo-iC3b beads were added to media wells (left) or confined agarose wells (right). The pHrodo-red signal from these merges has been isolated alongside the merge to emphasize internalized pHrodo-iC3b-bead signal. Scale bar represents 100µm. For all graphs, black points demonstrate experiment means, colored points demonstrate individual cell values for each run. N = 4 experiments for migration (A-E) or N = 3 experiments for phagocytosis (F-J); n = 50 cells for each condition per experiment for migration (A-C), n = 10 fields of view per condition for coating phagocytosis (E), or n = 15 fields of view for each condition per experiment for phagocytosis (F-J). Statistical analysis was assessed using via Kruskal–Wallis test with Dunn multiple comparisons: ns = not significant, *p < 0.05, **p < 0.01, ***p < 0.001, ****p < 0.0001. Error Bars represent SEM.

### The Arp2/3 complex is necessary for iC3b phagocytosis

The Arp2/3 complex has previously been implicated in iC3b-mediated phagocytosis (*17*), haptotaxis (*27*), and haptotaxis under confinement (*25*) in macrophage cells. We hypothesized that inhibition of the Arp2/3 complex would similarly impair BV2 phagocytosis and cell migration. Reutilizing our 4-chambered dish, we combined aspects from our previous two experiments to interrogate the Arp2/3 complex. We coated our 4-chamber dish with 10µg/mL AF-iC3b and then added agarose to half of the wells. We then also incorporated either DMSO vehicle control or the small molecular Arp2/3 complex inhibitor CK-666 into the media or agarose. Arp2/3 complex inhibition decreased migration velocity and accumulated distance compared to control, which was exacerbated further under confinement (**Figure 2A-B**).

Confinement alone did not shift these values, as both DMSO groups were similar to each other. However, confined DMSO-treated cells were more directionally persistent than unconfined DMSO-treated cells. (**Figure 2C**). Notably, CK-666 treatment lowered unconfined migratory persistence compared to DMSO, and confinement was unable to rescue the persistence phenotype in Arp2/3 complex-deficient cells (**Figure 2C**).

Interestingly, confinement does not significantly increase phagocytic uptake of surface-labeled AF-iC3b relative to unconfined cells in DMSO-treated conditions (**Figure 2D, visualized in 2E**). This is in contrast to confinement increasing phagocytic uptake of iC3b-labeled beads (**Figure 1F**). Arp2/3 complex inhibition decreased uptake of AF-iC3b compared to relevant DMSO-treated wells in unconfined and confined settings (**Figure 2D, visualized in 2E**). Confinement was unable to rescue AF-iC3b phagocytosis after inhibition of the Arp2/3 complex.

We wanted to then assess whether the Arp2/3 complex was involved in uptake of iC3b-beads. Therefore, we repeated our experiment with CK-666 and confinement, swapping out the AF-iC3b surface coating for pHrodo-iC3b-beads. We observed that confinement failed to increase phagocytosis in the presence of DMSO, pointing toward a slightly suppressive effect (**Figure 2F**) not observed when DMSO was absent (**Figure 1F**). DMSO interacts with sodium channels in CNS cells, prolonging pro-inflammatory onset of cells that could suppress phagocytosis (*28*, *29*). Nonetheless, CK-666 treatment disrupted phagocytosis in confined and unconfined settings when compared to DMSO control (**Figure 2F**). However, in this case confinement partially rescued the CK-666-mediated decrease in phagocytosis (**Figure 2F, visualized in Figure 2J**). Phagocytic impairment did not result in a change in phagosome size or number between DMSO and CK-666 treated cells assessed in the same confinement context (**Figure 2G-I**).

Interaction of cells with surface-labeled AF-iC3b and bead-opsonized pHrodo-iC3b resulted in lowered cell migration and cell phagocytosis upon Arp2/3 inhibition. Confinement was unable to rescue phagocytosis in iC3b coated on the dish bottom but was able to partially rescue phagocytosis when presented with iC3b opsonized to beads. These experiments using surface-labeled AF-iC3b led us to next examine whether iC3b is a haptotactic cue.

### Arp2/3 complex is required for iC3b haptotaxis but not confinement-induced migration on iC3b

Haptotaxis is directed, persistent cell migration in response to an increasing concentration of a substrate-bound cue. ECM gradients such as fibronectin are well-characterized haptotactic cues, (*30*), but it is unknown whether complement proteins like iC3b stimulate a directional migratory response. The effect of Arp2/3 complex inhibition previously demonstrated in the present study, combined with its known involvement in haptotaxis, points towards the potential for iC3b haptotaxis in synaptic pruning. As a first step to clarify this, we utilized the haptotaxis setup established previously in the lab to study fibronectin gradients (*25*). For the present study, we generated a gradient of AF-iC3b starting with a 250µg/mL source concentration in the central well (**Figure 3A**). Wells were spaced 1.25mm apart, and after an hour of allowing the gradient to establish, we seeded BV-2 cells into the outer wells and allowed them to migrate for 24 hours.

**Figure 3:**
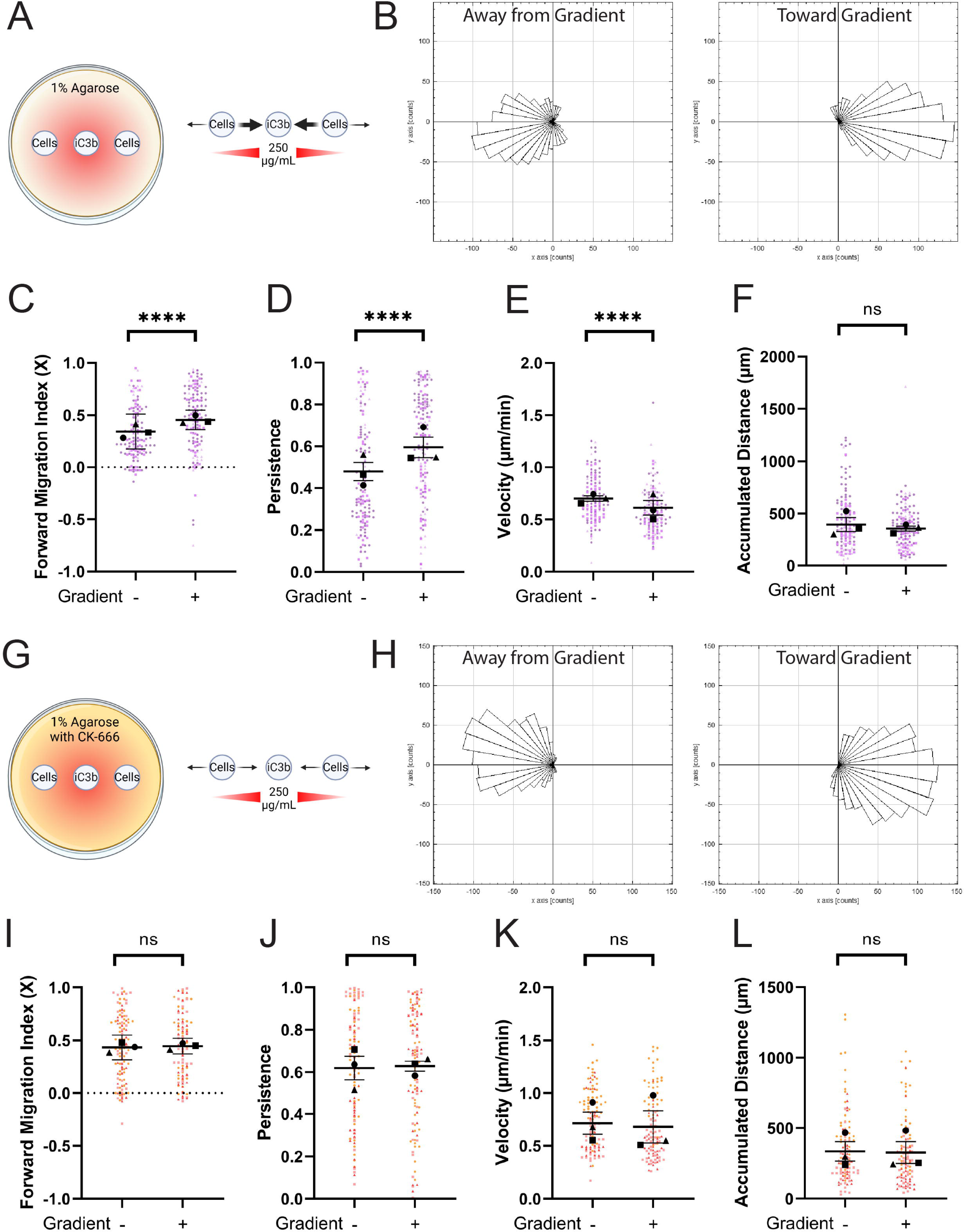
The Arp2/3 complex is required for haptotactic response to iC3b but not for directional confined migration on iC3b. A) Schematic depicting haptotaxis apparatus. Wells are placed 1.25mm apart. 250µg/mL AF-iC3b is placed in the center well and allowed to spread for 1 hour before washing. Cells are added to outer wells and then allowed to move under the agarose for 24 hours either towards the center well (toward gradient, denoted (+)) or away from the center well (away from gradient, denoted (-)). Strength of movement is noted in arrow thickness. B) example histograms of cell tracks demonstrating the number of cells traveling in each direction. C) Forward Migration Index in the X direction, measuring cell tracks in a binary system moving toward or away from the AF-iC3b haptotactic cue. D) Persistence of the cell during the length of its track. E) Velocity of cells migrating (µm/min). F) The maximum accumulated distance (µm) that the cell travelled. G) Schematic depicting haptotaxis apparatus with addition of 125µM CK-666 to agarose. Strength of cellular response to gradient cue is noted in arrow thickness. H-L) same type of graphs as in (B-F), noting now the addition of CK-666 to impacting cell motility. For all graphs, black points demonstrate experiment means, colored points demonstrate individual cell values for each run. N = 3 experiments for each graph; n = 50 cells for each direction per experiment for migration. Statistical analysis was assessed using the Mann–Whitney tests: ns = not significant, ****p < 0.0001. Error Bars represent SEM in all graphs except for FMIx, where Error Bars represent 95% Confidence Interval.

To ensure the gradient was present we measured from the inner edge of one cell well across the center well to the inner edge of the second cell well at the start and end of each run (**Supplemental Figure S3A-B**). These data confirm the consistency and stability of the iC3b gradient over time. Cell migration patterns were measured in what we previously established as “Zone 2”, one field of view displaced from the edge of the left and right cell wells (*25*). Notably, cells in each of the outer wells will respond by moving under agarose with the gradient (+, toward the center well) or against it (-, away from the center well). Thus, it is possible to directly compare the two populations of cells moving in the + and – directions within the same experiment. TIRF imaging of the glass bottom of our dishes confirmed that iC3b is bound to the glass and forms a gradient under the agarose (**Supplemental Figure S3C**).

BV-2 cells responded haptotactically to the iC3b gradient. Example histograms of cell motility demonstrate more cells moving along the X axis toward the gradient (+), while cells moving away from the gradient (-) have a more highly spread rose plot with fewer cells moving as strongly along the X axis (**Figure 3B**). The gradient in our experiments was established along the X-axis, so we quantified haptotaxis specifically via FMIx. In these experiments, FMIx was significantly increased when cells moved with the gradient, compared to away from it (**Figure 3C**), demonstrating the haptotactic influence of iC3b. Cell persistence mirrors the FMIx results, as cells moved with higher persistence toward the iC3b gradient (**Figure 3D**). Cells moving along the iC3b gradient (+) move with lower velocity compared to cells in the (-) orientation, but showed no change in total distance (**Figure 3E-F**).

The Arp2/3 complex is a critical effector of haptotactic sensing, as previously demonstrated in macrophages responding to fibronectin (*25*, *27*). To assay this in BV-2 cells, we repeated our experiment with CK-666 polymerized into the agarose (**Figure 3G, Supplemental Figure S3D-E**). When exposed to CK-666, BV-2 cells moving along the iC3b gradient had a more highly spread rose plot than untreated cells moving under the same conditions (compare ‘toward gradient’ populations in **Figure 3H** and **Figure 3B**), similar to cells moving against the gradient in both datasets. FMIx quantification demonstrates this further, as there is no statistical difference between cells moving toward (+) or away (-) from the gradient when CK-666 is present (**Figure 3I**). While untreated cells increase migratory persistence and decrease velocity on an iC3b gradient (+ population) (**Figure 3D**), both measures are similar in (+) and (-) populations when CK-666 is present (**Figure 3J-K**). While Arp2/3 is required for directional migration under these conditions, Arp2/3-deficient cells are capable of moving well under agarose. As with the untreated cells, CK-666 had no effect on distance traveled during migration (**Figure 3L**). These data together demonstrate the haptotactic nature of iC3b, and suggest that Arp2/3-generated lamellipodial protrusions are necessary for microglia to detect iC3b gradients.

### The Arp2/3 complex maintains microglial morphology *in situ*

We then had the unique opportunity to translate our *in vitro* results into a system with more *in vivo* relevance: live imaging of microglia *in situ* in hippocampal slice cultures. Utilizing CX3CR1^+/GFP^ mice (*31*), we created 400µm coronal slices of mouse brains from mice between 35 and 50 days old, of equal sex representation and imaged in the CA1 region of the hippocampus. To examine the effects of Arp2/3 complex inhibition, slices were incubated in either 200 µM CK-666 or DMSO vehicle control prior to imaging. 200 µM CK-666 concentration was chosen based on previous literature utilizing CK-666 in hippocampal slices (*32*).

We started by examining microglial surveillance. Individual cells were imaged every minute for 10 minutes to examine dynamic movement under our different treatment groups: no treatment (negative control), DMSO (vehicle control), or 200 µM CK-666 (**skeletons visualized in Figure 4A**). First, we measured the morphological changes in cells when treated with either DMSO or CK-666 at the first time point in our surveillance videos. CK-666 treatment significantly decreased the number of branches, junction points, and process tips compared to both DMSO-treated and non-treated cells (**Figure 4B**). These data indicate that Arp2/3 complex plays a major role in maintaining microglial morphology *in situ*. There was no statistically significant difference between the DMSO and non-treated cells.

**Figure 4:**
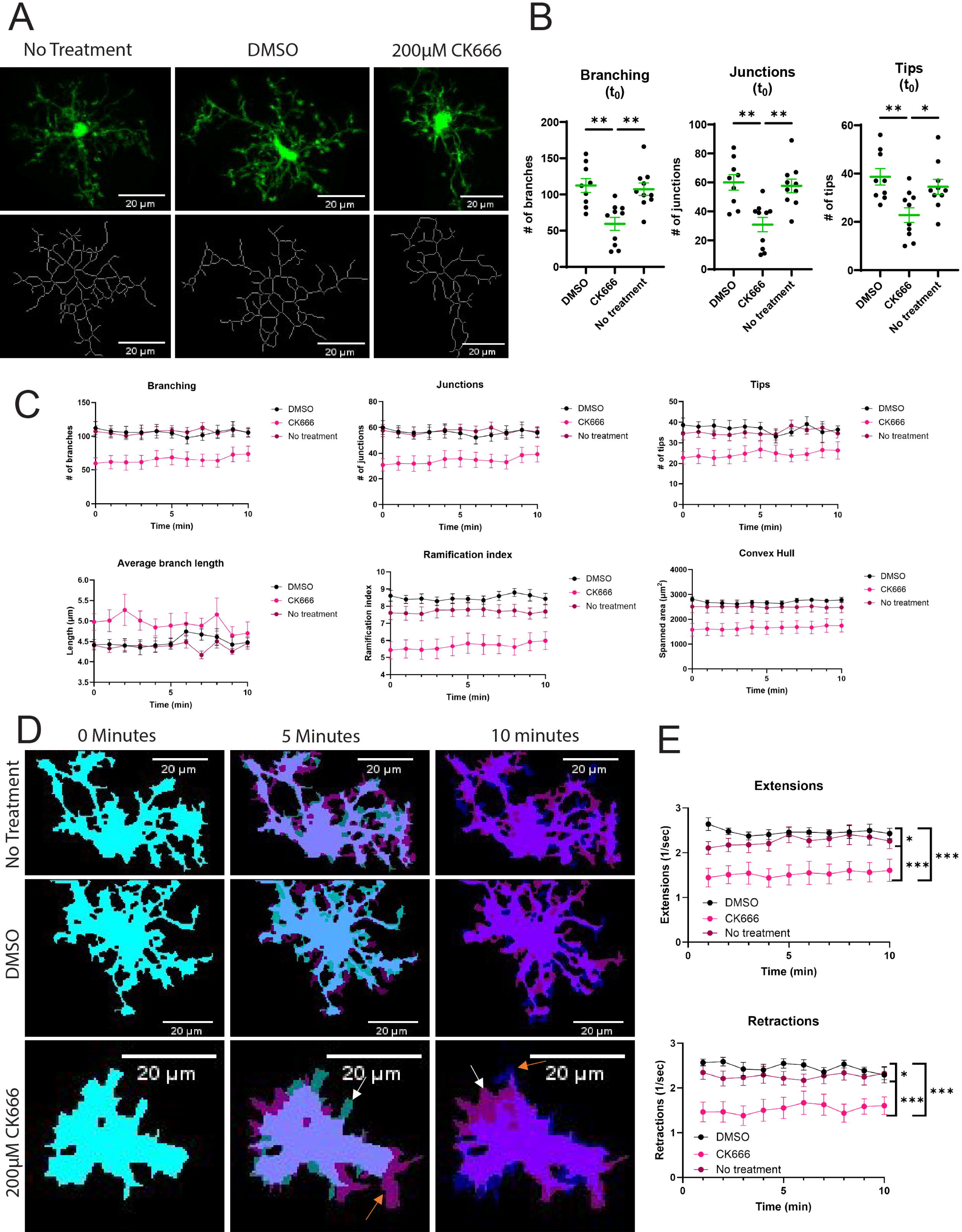
Morphological features of hippocampal slice microglia treated with CK666, vehicle control (DMSO), or untreated. A) Original images (top) with skeletons generated using MotiQ 2D Analyzer (bottom) at first imaging timepoint after drug washout. Scale bar represents 20µm. B) Quantification of branching, junctions, and tips at first imaging timepoint (n=9-10 cells). Statistical analysis performed using one-way ANOVA (* *p*-value < 0.05, ** < 0.01), lines and error bars represent mean ± SEM. C) Minute-to-minute morphological features after drug washout (mean ± SEM). D) Binarized image with ‘mean’ algorithm and closed function. Colorized masks represent independent timepoints (0 min = cyan, 5 min = magenta, 10 min = blue). Non-overlapping regions represent areas of extensions and retractions. White arrows pointing examples of retraction; orange arrows pointing examples of extension. Scale bar represents 20µm. E) MotiQ automated quantification of extensions and retractions (mean ± SEM). Statistical analysis performed using an ordinary 2-way ANOVA with Tukey’s multiple comparisons test of treatments. Main effect of drug treatment (F(2, 258) = 74.71, p<0.0001***) with no time-dependent effect or interaction. Between group comparisons were significant (* p-value<0.05*, <0.01**, <0.0001***).

We then examined changes in cell dynamics over the course of our videos. Overall, we saw similar results in our non-treated and our DMSO-treated cells, with CK-666 cells demonstrating significantly different values in many morphology measurements (**Figure 4C**). As with our single timepoint analysis, CK-666 treatment caused microglia to have fewer branches, fewer junctions, and fewer tips along their existing branches over the entire time course (**Figure 4C**). Overall processes are limited as well in CK-666 treated microglia, reflected by lower ramification index (overall complexity of the microglial skeleton) and a smaller convex hull measurement (spanned area of the cells) as the cells are taking less area within their 3D environment compared to untreated and DMSO-treated counterparts (**Figure 4C**).

We also examined dynamic protrusion of our cells to test whether surveillance-like behaviors are impacted by Arp2/3 disruption. Indeed, CK-666 treatment impairs protrusion (extensions) and retraction compared to controls (**Figure 4D, quantified in Figure 4E**). Interestingly, DMSO treatment increased protrusion (extensions) and retraction compared to untreated counterparts, suggesting that vehicle treatment elicits a somewhat more dynamic surveillance behavior in these cells (**Figure 4E**). Given the profound contribution of Arp2/3 to maintaining microglial morphology and supporting surveillance-related cell protrusion, we wanted to next evaluate how these alterations in cellular structure and behavior would impact microenvironmental sensing of chemotactic and haptotactic cues.

### Arp2/3 complex is required for iC3b sensing, but not ATP-induced chemotaxis

The Arp2/3 complex is necessary for macrophage haptotaxis but not chemotaxis (*27*). However, the differential contribution of Arp2/3 to specific extracellular sensing pathways has not been as robustly studied *in vivo* in mice. Our hippocampal slice culture presents an ideal opportunity to evaluate this question using endogenous microglia in a physiologically relevant setting *in situ*. We decided to use ATP as a chemotactic cue, due to its established history as a microglial chemotactic cue (*33–35*). Therefore, we developed a system to inject vehicle (tissue bath solution) or 1mM ATP into DMSO- or CK-666-treated brain slice cultures using a micropipette.

We imaged brain slices for 1 hour at 2-minute intervals, with ATP injection at 8 minutes, and continued imaging for 52 minutes. After the ATP injection, a contracting ring of microglial processes extended toward the pipette end until the pipette was blocked, resulting in two occurrences: 1) one or multiple processes climbed the inside of the pipette to further respond to the ATP, and 2) the external processes that had been responding stopped, as there was no longer a source of ATP disseminating. DMSO- and CK-666-treated cells responded similarly to ATP. Microglia did not respond to injection of tissue bath solution, while ATP injection elicited a much shorter response time that was not affected by Arp2/3 inhibition (**Figure 5A**). Likewise, ATP elicited process generation equally from DMSO- and CK-666-treated microglia (**Figure 5B**). Finally, both DMSO- and CK-666-treated cells were equally capable of crawling up the ATP-containing pipette until they clogged it (**Figure 5C**). When mice were separated by sex, we only noted a faster response time in our male mice for CK-666 treated microglia (**Figure 5D**), while none of the other measurements were significant (**Figure 5E-F**). There was also no significant difference between male and female values within DMSO or within CK-666 treatments. In total, we observed no meaningful difference in microglial response to ATP upon Arp2/3 disruption. These data suggest that microglial chemotactic sensing *in vivo* is not likely to be impaired by loss of Arp2/3 function.

**Figure 5:**
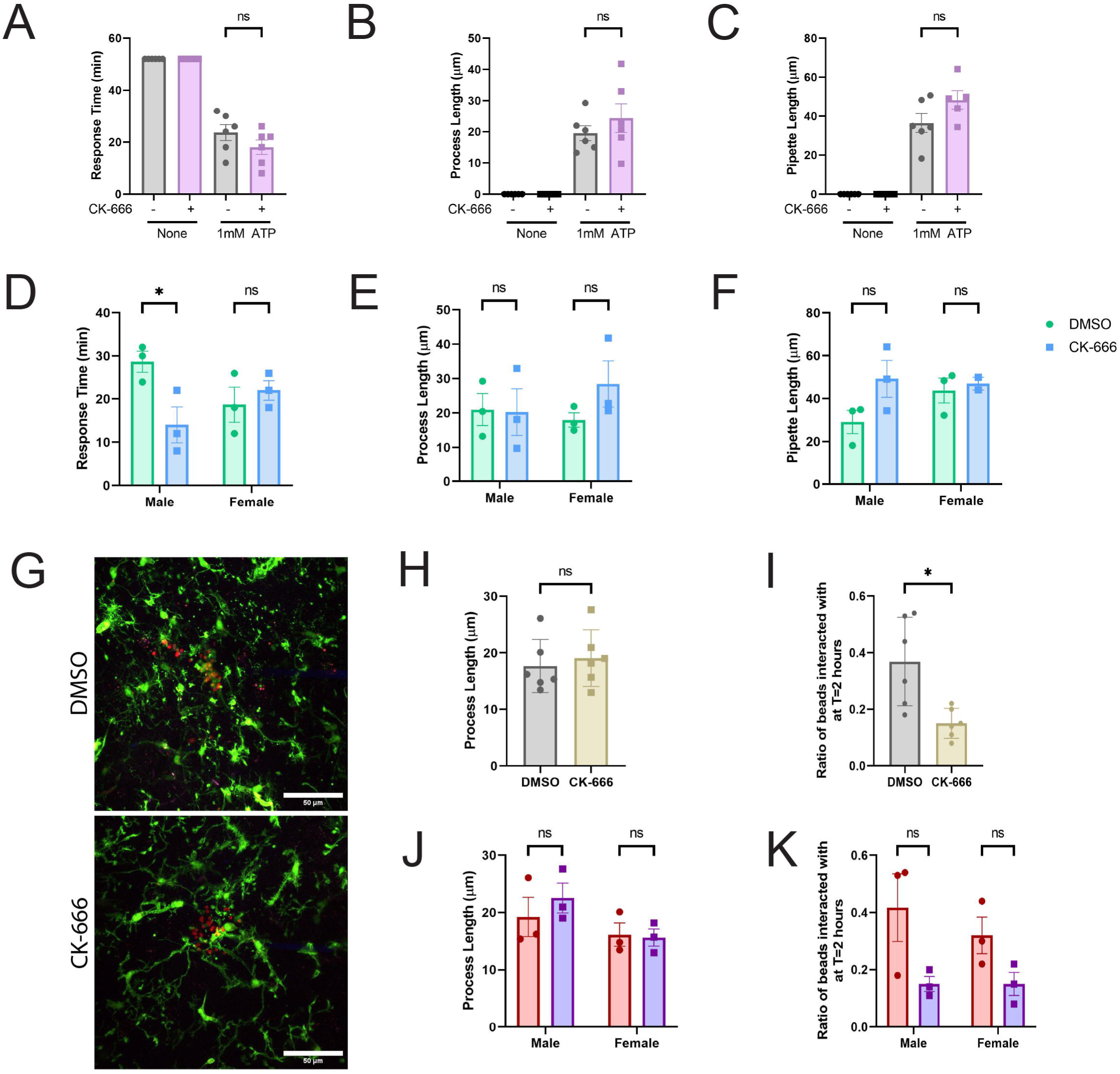
The Arp2/3 complex is required for cellular response to iC3b but not ATP-dependent chemotaxis *in situ*. A) Measured response time from cells receiving chemotactic cue of 1mM ATP to the processes blocking the pipette, in minutes. B) Averaged process length (µm) from cells responding to chemotactic cue. C) Length (in µm) of micropipette that a process has visibly crawled up by end of 1 hour video. D-F) ATP results of (A-C) broken down into male and female specific results to examine sex differences. G) Example endpoint images of 2 hour phagocytosis assay. Scale bar represents 50µm. H) Averaged process length (µm) from cells responding to pHrodo-iC3b beads cue. I) Ratio of beads interacted with vs not interacted with at the end of the 2 hour assay. J-K) Results from (H-I) broken down into male and female mice to examine sex differences. N = 6 experiments for each graph (3 male; 3 female). Statistical analysis was assessed using the Mann–Whitney tests (Non-Sex graphs): ns = not significant, *p < 0.05. Statistical analysis was assessed using 2way ANOVA and multiple t tests for graphs examining the impact of sex: ns = not significant, *p < 0.05. Error Bars represent SEM in all graphs.

Next, we adapted the micropipette injection protocol to incorporate pHrodo-iC3b beads to examine whether microglia demonstrate an Arp2/3-dependent haptotactic response *in situ*. In these experiments, hippocampal slices were imaged for 2 hours with 2-minute intervals. pHrodo-iC3b beads were injected at the 8-minute mark, similarly to ATP injection (**Figure 5G**).

Interestingly, process lengths were not altered by Arp2/3 disruption (**Fig. 5H**), even though microglia interaction with iC3b beads was dramatically reduced by CK-666 treatment (**Fig. 5I**). Unlike with ATP treatment, iC3b beads did not elicit process extensions toward the iC3b (**Compare 5B to 5I**). These data suggest that the response to iC3b-labeled targets is logically a more ‘local’ phenomenon than chemotaxis, as microglia are likely contacting iC3b labeled spines *in vivo* via local surveillance. We did not observe a sex difference in how microglia responded to iC3b (**Figure 5J-5K**). Local iC3b gradients could help recruit microglia to areas where they are required for spine pruning without requiring a large response from the regional population. Together our data suggests that microglia haptotaxis is Arp2/3-dependent in an *in vivo*-like environment.

## DISCUSSION

We set out to examine the Arp2/3 complex’s contribution to microglial function, and to determine the nature of iC3b sensing *in vitro* and *in situ*. BV-2 cells move less on iC3b than on fibronectin, likely due to their propensity to phagocytose iC3b from the bottom of the dish in an Arp2/3-dependent fashion. With iC3b-opsonized beads, phagocytosis increased under confinement, again in an Arp2/3 complex-dependent fashion. Likewise, when these cells were tasked with moving under confinement on iC3b in the presence of CK-666, we see a decrease in migratory abilities including loss of haptotaxis and a decrease in phagocytosis that is not rescued by confinement. Finally, we translated these findings to *in situ* studies of microglia in brain slices. While Arp2/3 complex is responsible for baseline surveillance and maintaining ramified microglia morphology, the chemotactic response of microglia processes to ATP was maintained in CK-666 treated slices. Conversely, CK-666-treated microglia could not extend stable processes that maintain contact with iC3b-labeled beads *in situ*, compared to vehicle-treated microglia. Together this body of work establishes that Arp2/3 complex-dependent haptosensing is an important feature of the microglial response to iC3b.

Though haptotaxis has been interrogated at the molecular level *in vitro*, much less is known about how haptosensing contributes to physiological functions. Leukocyte haptotaxis is perhaps better contextualized *in vivo* than for other cell types. One striking example is dendritic cells’ ability to haptotax along a gradient of immobilized chemokine CCL21, with compelling evidence pointing to *in vivo* CCL21 gradients (*36*). Within the CNS, axonal growth cones of newly developing neurons in both the brain and spinal cord follow gradients of both chemoattractant and haptoattractant cues, with new studies pointing towards netrin and sonic hedgehog as haptotactic cues for neurons (*37*, *38*). Recent studies have found that calvarial osteoblast progenitors haptotax along the embryonic cranial mesenchyme (*39*). Conversely, disruption of haptosening can be detrimental. Tumor cell invasion has been shown to downregulate actin cytoskeleton components responsible for haptotactic sensing to enable this migration (*40*, *41*). Our work contributes to this burgeoning area by demonstrating that iC3b elicits a haptotactic response, and that disrupting microglial haptosensing in the context of a normal CNS microenvironment impairs the response to complement-opsonized ligands. The role of the Arp2/3 complex in integrin-dependent phagocytosis and motility has been established *in vitro* (*17*). Conversely, numerous studies in multiple cell types confirm that Arp2/3 is dispensable for chemotaxis (*17*, *27*, *42–44*). However, this dichotomy has not been thoroughly explored in mammalian systems *in vivo*. In the present study we demonstrated that Arp2/3 complex-deficient microglia in their normal *in situ* environment respond dramatically to a chemotactic ligand while failing to interact with local iC3b cues. These findings suggest that cells are capable of integrating cues from their environment using distinct sensing pathways, and that cytoskeletal responses to the microenvironment may be much more specific than previously suspected.

Two different models of synaptic pruning have been proposed: direct trogocytosis via microglial interaction with dendritic spines, or neural shedding wherein the spine detaches from the neuron and microglia phagocytose these shed spines within the ECM (*45*). The neural shedding model is highly reminiscent of a haptotactic gradient. iC3b as a haptotactic ligand would point towards a better explanation for its use in inflammatory immune responses and synaptic pruning (*46*). iC3b could act equally well as a local haptotactic cue in either case. Future studies are important in further clarifying how iC3b-dependent haptosensing contributes to microglial synaptic pruning.

Trogocytosis, is an important aspect of environmental surveillance that could be further assessed by our *in vitro* surface coating assays. During trogocytosis phagocytes pinch off pieces of a cell rather than fully ingesting them (*22*). Studies in synaptic pruning support the idea that microglial synaptic pruning of the presynaptic synaptic bulb at the axon terminal occurs via trogocytosis, both in hippocampal slices *in situ* (*47*) and in amphibians *in vivo* (*48*). But these studies have not focused on the means of clearance of the postsynaptic side of the cell. While previous studies have been firm in labelling this as complement-mediated phagocytosis (*11*, *49*), iC3b surface coating assays allow for the visualization of iC3b uptake via a process potentially similar to trogocytosis. The Arp2/3 complex, which we confirm is required for iC3b uptake, creates part of the cell tunnel involved in forming the trogocytic cup (*23*). Therefore, our approach may be a first step toward modeling synaptic pruning more effectively *in vitro*.

Neurodegenerative diseases such as Alzheimer’s and Parkinson’s disease both present with chronic neuroinflammation. Evidence points toward activated microglia phagocytizing dendritic spines prematurely and contributing to cognitive decline in these diseases due to hastened synapse loss (*50*). Multiple neurological disease model mouse studies have found that ablation of C1q (another complement component), C3, or CR3 improved learning and memory tasks (*4*, *51*– *54*). Microglial ablation has also improved learning and memory tasks in neurological disease model mice as well (*55*, *56*). These studies suggest a need to mitigate hyperactive synaptic pruning in neurodegenerative disease models. Therefore, targeting the microglial haptosensing pathway therapeutically might preserve synapses and ameliorate neurodegenerative disease.

The Arp2/3 complex serves as a proof of concept for this idea. Inhibition of the Arp2/3 complex selectively targets haptosensing of iC3b, but preserves microglial chemotaxis and process generation in response to ATP. Previous research in macrophages demonstrated that Arp2/3 complex is necessary in iC3b but not IgG phagocytosis (*17*), once more pointing toward specific effects of Arp2/3 disruption. Prior *in vivo* research involving the Arp2/3 complex centers around the impact of its loss in neurons on dendritic spine formation (*21*). To our knowledge, genetic targeting of the Arp2/3 complex within microglia has not yet been demonstrated *in vivo* in mice. A potential confounding factor in doing is related to challenges utilizing Cre to target microglia *in vivo*, including leaky Cre drivers, potential for non-cell specific activation, and potential cytotoxicity of some Cre-targeting lines (*57–59*). Considering Kim *et al*.’s study of Arp2/3 loss in neurons (*21*), utilizing more general approaches could have side effects beyond targeting synaptic pruning. Our findings suggest that targeting iC3b haptosensing as specifically as possible in microglia may effectively pause synaptic pruning and slow neurodegeneration *in vivo*.

## MATERIALS AND METHODS

### Cells

Established murine microglial cell line BV-2s were used for all experiments listed. Cells were grown in complete cell culture media, containing Dulbecco’s Modified Eagle Medium (DMEM) (Gibco, 31053028), 5% fetal bovine serum (FBS) (Gibco, 10437028), 1% glutaMAX (Gibco, 35050061), and 1x Antibacterial-Antimycotic (Gibco, 15240062) at 37°C and 5% CO_2_. 70-80% confluent dishes were treated with 0.05% Trypsin-EDTA solution (Caisson Labs TRL02-100ML) at 37°C for 10 minutes. The solution was then aspirated and replaced with 2mL of complete cell culture media that was gently sprayed over the bottom of the dish to dislodge cells and collect for counting and passage into new dishes.

### Labelling iC3b with Alexa Fluor

Alexa Fluor 555 Fluorescent Protein Labeling Kit (Thermo Fisher, A20174) was used as directed to label 250µM of Human iC3b (CompTech, A115).

Western Blot analysis was performed with the Alexa Fluor 555 labeled iC3b (AF-iC3b), utilizing a Ponceau Stain to verify that the iC3b is labeled and there was no excess dye present in the solution.

### Adhesion assay

8 well chamber dishes (Cellvis, C8-1.5H-N) were prepared with a coating of 10μg/mL fibronectin (Gibco, 33016015) or AF-iC3b as previously described, then all wells were filled with PBS. Confluent BV2 dishes were handled and cells counted as previously described. 0.5mL Eppendorf tubes were each filled with 100µL of complete cell culture media and 40,000 cells. Tubes were spun down at 1000xg for 5 minutes, then media aspirated. Cells were resuspended in 47.5µL of 2% FBS (diluted in PBS) and 2.5µL of the desired 1mg/mL antibody to create a 50µg/mL antibody concentration. Antibodies used were Normal Rat IgG (Invitrogen, 31933), Cd11b (Abcam, ab8878), and Integrin β2 (NovusBio, NBP1-41272). Tubes were placed at 37°C for 10 minutes. Cells were spun down again and resuspended in 40µL of complete cell culture media. The 8 well chamber dish was then prepped with 1mL of media in each well and half of the cells transferred from each antibody tube to the corresponding well. Dishes were placed on a BioTek Cytation microscope, incubated at 37°C, and imaged every 30 minutes for 2 hours. Plates were then removed and received one wash of 1x Cell Culture Phosphate Buffered Saline (Corning, MT21040CV) (referred to from here on as PBS) before placed back on the microscope and a final image was taken.

### Analysis of adhesion assay

Image files were opened in ImageJ. Using the Counting Cells plugin, the total cells per field of view both pre- and post-wash were counted. Counts were transferred into GraphPad to graph. Graphs represent the percentage of pre-cells remaining, the percentage of lost cells in the wash, and the total number of cells present during the pre-wash stage.

### 4 chamber dish creation (iC3b coating phagocytosis)

4-chamber well dishes were coated with fibronectin (10μg/mL) or AF-iC3b (1μg/mL, 10μg/mL, or 20μg/mL) for 1 hour at 37°C. Chambered dishes were washed three times with PBS. Two wells were each filled with 1mL PBS and two wells were each filled with 1mL 1% agarose as previously described.

### Motility microscope assay

Upon completing the 3 hour incubation, chambered dishes were placed in an environmental chamber on a Tokai Hit INU incubation system controller on the Olympus IX83 to be held at 37°C, 90% humidity, and 5% CO_2_ for the duration of the live cell imaging. A 20X air objective was employed during overnight time lapse imaging. 10 fields of view per well were selected. Images were taken at each field of view in the relief contrast channel every 10 minutes for the span of 16 hours.

### Analyzing motility

Each motility file was opened in FIJI ImageJ. The Manual Tracking plugin was used to track 10 cells per field of view and combining data across all fields of view within treatment groups. Cell tracking was stopped if a cell divided or ran into another cell, resulting in a changed direction. The resulting measurements were uploaded into the Ibidi Chemotaxis and Migration Tool ImageJ plugin to measure velocity, accumulated distance, and persistence.

Persistence (also termed d/T) is a measure of the straight-line distance between a cell’s starting and ending position on a track (d) divided by its overall track length (T). Thus, a value very close to zero (high T) is understood to be meandering much more than a cell moving along the shortest distance between its start and end (d = T), which has a value of 1. The closer the value to 1, the more persistent the migration is. These data tables were exported to GraphPad Prism to graph differences in velocity, distance, and persistence between treatment groups.

### Analysis of iC3b eating

Experimental images for the three AF-iC3b wells were loaded into ImageJ and the relief contrast channel removed. The BioVoxxel Pseudo Flat Field Correction plugin was used to remove uneven fluorescence from the Olympus IX83 capturing ability. The image was made binary and then the raw intensity difference was measured for the first and last time point in the stack. The change in raw intensity between the two timepoints quantified how much of the iC3b fluorescence was no longer visible, signifying uptake by the cells.

### 4 chamber dish creation (bead phagocytosis)

4 well chamber dishes with #1.5 coverglass bottoms (Cellvis, C4-1.5H-N) were coated in 10μg/mL fibronectin (Gibco, 33016015) for 1 hour at 37°C. Chambered dishes were washed three times with PBS. Two wells were each filled with 1mL PBS and two wells were each filled with 1mL 1% agarose. Agarose was prepared as follows. 2.5mL of 2x Hank’s Balanced Salt Solution (Sigma, H1387-10X1L) and 5mL of complete cell culture media (as defined above) were combined and placed in a 68°C bath for 1 hour. 0.12g of low gelling temperature agarose (Sigma, A9045-5G) was added to 2.5mL of sterile water (Corning, 25-055-CV) and heated in a microwave until fully dissolved. The two solutions were combined to create a warm 1% agarose mixture. 1mL of said mixture was poured into each well and allowed to solidify for 90 minutes at room temperature. Chambered dishes were wrapped in parafilm and stored at 4°C for up to one month.

### Phagocytosis bead creation

Label 100µg Human iC3b (CompTech, A115) using pHrodo™ iFL Red Microscale Protein Labeling Kit (ThermoFisher, P36014) according to manufacturer instructions. Precipitated labeled protein was opsonized to 30µL of 2-micron Polybead Carboxylate Microspheres (Polysciences, Inc, 18327-10) (hereafter referred to as beads) in 1mL of 1x PBS at 4°C overnight. Beads received three washes with 1x PBS after completing opsonization and were resuspended in a final volume of 500µL, stored at 4°C for up to 4 months. Before usage in an experiment, beads were vortexed vigorously.

### Cell preparation

Agarose chambered dishes were prepped for cells by replacing PBS with 1mL complete cell culture media. Agarose wells had two punches placed in each for cell and bead insertion using a 0.75mm biopsy punch tool (LabTech, 52-004908). Confluent dishes were treated according to the standard passaging protocol (above). For cell migration experiments, 13,000 cells were added to each media well. For phagocytosis, 18,000 cells were added to each media well. Lower numbers were used for migration experiments to minimize cell-cell interactions and allow more room for random migration uninterrupted. For both types of experiments in the confined condition, 15,000 cells were inserted under the agarose in each punched location using a gel-loading pipette tip. If the amount of media containing 15,000 cells exceeded 15µL, the cells were spun down at 1000xg and resuspended in 10µL of complete cell culture media. Chambered dishes were placed at 37°C for three hours to allow for cells to sit down and spread before being moved to the microscope. For the inhibitor trials, each cell culture media well received either vehicle or inhibitor to create the working concentration used for cell treatment (more information below).

### Phagocytosis microscope assay

Upon completing the 3-hour incubation, iC3b beads were added to wells. Media wells received 5µL of beads. For confined wells, a 1:4 dilution of the beads was made with PBS and inserted into each punch in the agarose to match bead density with the media condition. Chambered dishes were then placed in a Tokai Hit INU incubation system controller on the Olympus IX83 to be held at 37°C, 90% humidity, and 5% CO_2_ for the duration of the live cell imaging. A 20X air objective was employed during overnight time lapse imaging. 15 fields of view per well were selected. Images were taken at each field of view every 10 minutes for the span of 8 hours. Both the relief contrast and the DsRed channels were utilized. The DsRed channel was set to 500ms exposure to detect pHrodo signal upon bead internalization by cells.

### Analyzing bead density

Each phagocytosis file was opened in Fiji ImageJ. A bandpass filter was then run on the relief contrast channel. Structures were filtered to fit between 5 and 15 pixels and then threshold filtered until only the beads were highlighted in red. The analyze particles function was used to count bead density. Area threshold was set to 10-30 micron^2^, circularity threshold was set to 0.8-1.0, and outlines were shown. When summarized, average size was maintained between 15 and 17 micron^2^ across the time lapse, with threshold being adjusted and analysis rerun if the averages were too small or too large. Counts of beads per time point were averaged to create the bead density designator for each file. Averaged beads numbers falling between 50 and 500 were designated low bead density; between 500 and 950 medium bead density; between 950 and 1550 high bead density.

### Analyzing phagocytic rate

Each phagocytosis file was opened in FIJI ImageJ. Channel colors were corrected and brightness/contrast adjusted to create the same brightness and contrast parameters for each file. Using the cell counter plugin, the total number of fluorescent and non-fluorescent cells was counted at each measured time point throughout the phagocytosis videos. Comparisons between treatment groups at individual time points as well as comparing changes over time within each group were conducted with GraphPad Prism. Graphs were normalized to T=0. Fluorescent cells at the beginning of the videos were removed from the count of both fluorescent cells and total cells per field of view. Data comparisons were broken up by bead density, only comparing media and confined fields in the low bead density category to each other.

### Analyzing phagosome size

Each phagocytosis file was opened in CellSens Dimension software under the Count and Measure plugin. Under detection options, minimum object size was set to 5 pixels. Under the Thresholding tab, the adaptive threshold was selected. Selecting the DsRed channel, the maximum was set to 65000 (the maximum) and the minimum was changed for each image to label only internalized beads, although never going below 1000. Selecting the relief contrast channel, the maximum was set to 2 and the minimum to 1 so that only fluorescent labels were detected. Outputs of total number of phagosomes and their average size per field of view were then verified at each timepoint before the data was exported to an excel file. This data was presented three ways. The average size of phagosome was reported by dividing the average size per field of view at 2 hour by the total number of phagosomes detected at 2 hour. Average size of phagosome per phagocytic cells was reported by dividing the average size per field of view by the number of phagocytic cells at that timepoint. The average number of phagosomes per phagocytic cell was reported by dividing the total number of phagosomes at 2 hours by the number of phagocytic cells at that timepoint. Average size of phagosome and average size of phagosome per phagocytic cell differ by the earlier examining if beads are trafficked into the same or different lysosomes and the latter examining if overall more beads are being trafficked in each treatment condition.

### 4 chamber dish creation (iC3b coating with CK-666 phagocytosis)

The small molecule Arp2/3 complex inhibitor CK-666 (Abcam, ab141231) was resuspended in anhydrous DMSO (ThermoFisher, D12345) to create a stock solution. 4-chamber well dishes were coated with 10μg/mL of AF-iC3b for 1 hour at 37°C. Chambered dishes were washed three times with PBS. Agarose was prepared as previously described and then split into two aliquots of 5mL warm agarose solution. Each aliquot received either vehicle (anhydrous DMSO) or CK-666 at a final concentration of 125μM. AF-iC3b coated dishes now contained two PBS wells, one vehicle agarose well, and one CK-666 agarose well. Chambered dishes were wrapped in parafilm and stored at 4°C for up to one month. Cells were seeded, incubated, imaged, and analyzed similarly to non-CK-666 treated runs.

### 4 chamber dish creation (bead phagocytosis with CK-666)

Agarose containing either DMSO or 125μM CK-666 was prepared as previously described. Agarose chambered dishes were prepared as previously described with 10µg/mL FN, now created with two PBS wells, one vehicle agarose well, and one drug agarose well. Chambered dishes were wrapped in parafilm and stored at 4°C for up to one month. Cells were seeded, incubated, given pHrodo-iC3b beads, imaged, and analyzed similarly to non-CK-666 treated runs.

### Haptotaxis Agarose Plate Preparation

1% agarose was made as previously described. 3mL of this mixture was poured into a 35mm glass bottom dish containing a 20mm micro-well (Cellvis, D35-20-1.5-N) and allowed to solidify for 90 minutes at room temperature. The plates were not pre-coated with substrate. Plates were wrapped in parafilm and stored at 4°C for up to one month, after which point the gel’s integrity could be compromised.

### Haptotaxis Cell and Dish Preparation

A 3.5mm biopsy punch tool (Robbins Instruments, RBP-35) was used to generate wells in the agarose gel. Punches were created and then excess gel was aspirated to create each well. A center well was created and AF-iC3b was allowed to spread under agarose from the central well for 1 hour at 37 °C. The well was washed once with PBS and then two linear wells of 1.25mm well spacing were punched in the agarose. Cells were prepared and counted as previously described. Once counted, 100,000 cells were transferred to 2 separate eppendorf tubes (200,000 cells total) and spun down at 1000xg for 3.5 minutes. Excess media was aspirated, and cells were resuspended in 20 µL of cell culture media. These suspensions were then transferred to each of the cell wells in the prepared agarose dish.

### Haptotaxis migration imaging

Cell plates were placed on the Olympus IX83, and an INU incubation system controller from Tokai Hit was used to maintain cells in a stable environment at 37°C, 90% humidity, and 5% CO2. Under-agarose chambers were placed into the environmental chamber. A 20X air objective was employed during overnight time lapse imaging. Cells were then imaged via relief contrast and the DsRed channel. A total of 32 positions were selected to assay macrophage haptotaxis for each experiment: 4 non-overlapping sites from the top to the bottom of the edge of a cell well, which composed Zone 1, 4 more non-overlapping sites at the same Y position but shifted approximately 665 μm (1 field of view) away, which composed Zone 2. This process was repeated on the other side of the same well, and then on both sides of the second cell well. An additional 12 images were taken across the length of the chamber between the (+) regions of the cell wells for fluorescence intensity analysis. Images were captured at 10-minute intervals for 24 hours.

### Haptotaxis migration analysis

Each of the Zone 2 files were opened in ImageJ. The Manual Tracking plugin was used to track each cell per field of view and combining data across fields of view within that file’s Zone 2 set of 4 files. Cell tracking was stopped if a cell divided or ran into another cell, resulting in a changed direction. The resulting measurements were uploaded into the Ibidi Chemotaxis and Migration Tool ImageJ plugin to measure track velocity, accumulated distance, persistence, and forward migration index in the X coordinate (FMIx). FMIx is calculated by dividing the difference between the starting and ending X coordinate of a cell track value by the accumulated distance. These data tables were exported to GraphPad Prism to graph differences in velocity, distance, and persistence between treatment groups. Note that persistence is not the same as FMIx as the persistence values do not consider any external frame of reference (e.g. an external gradient).

### Alexa Fluor 555 iC3b (AF-iC3b) Quantification

AF-iC3b was allowed to spread under agarose from the central well for 1 hour at 37°C prior to each migration assay. 12 images were taken across the length of the chamber between the (+) regions of the cell wells during migration runs. Each position was opened in ImageJ, and the DsRed channel was isolated. A 500W x 1H pixel line is created through the center of the image to measure the fluorescence intensity. A Gray Value plot was generated, and the data was saved to Excel and then loaded into Graphpad Prism.

### CK-666 Under-Agarose Haptotaxis Assay

Agarose was prepared as previously described, with the addition of 125µM CK-666 to the final gel volume. 3mL of this mixture was added to each 35mm dish. The agarose chamber was then prepared as previously described with one source well and two cell wells. 20µL of 250µg/mL AF-iC3b was added to the source well and allowed to diffuse for 1 hour at 37°C. After 1 hour, the source well was washed once with PBS. Cells were prepared as previously described and seeded into the plate. Imaging and analysis was conducted as previously described.

### Hippocampal slice preparation

CX3CR1^+/GFP^ mice were housed according to university and IACUC standards. Equal numbers of male and female mice were used in these experiments. Between 35 and 50 days of age, mice were sacrificed by CO2 inhalation. 400-μm thick coronal sections were cut in 4°C dissection solution (high-sucrose artificial cerebrospinal fluid (aCSF) containing (in mM): KCl, 2; NaH2PO4·H2O, 1.25; MgSO4·7H2O, 2; MgCl·6H2O, 1; NaHCO3, 26; CaCl2·2H2O, 1; D-glucose, 10; sucrose, 206; bubbled with a mixture of 95% O2 / 5% CO2) on a Leica VT1200S Vibratome (Buffalo Grove, IL, USA) and subsequently incubated for 30 minutes at 37°C in normal aCSF (containing (in mM): NaCl, 126; KCl, 3, NaH2PO4·H2O, 1.25; MgSO4·7H2O, 2; NaHCO3, 26; CaCl2·2H2O, 2; D-glucose, 10; sucrose, 20; bubbled with a mixture of 95% O2 / 5% CO2).

Brain slices were kept at room temperature for up to 8 hours. For imaging, the slices were mounted into a tissue slice chamber (Warner Instruments) and perfused with normal aCSF at room temperature. A ZEISS 7MP microscope was used to perform 2-photon fluorescence microscopy approximately 80 μm below the surface of the slice. The microscope system used a pulsed infrared laser (Chameleon Vision2, Coherent, Santa Clara, CA) with a tunable wavelength range from 680-1080 nm and was set to 850 nm for the time series experiments and 800 nm for the overview images respectively. Fluorescence was detected with three non-descanned detectors (NDDs) equipped with a 480 nm short pass for blue detection (dextran), a 525/50 nm bandpass for green detection (GFP) and a 605/70nm bandpass for red detection (pHrodo-iC3b labeled beads). Imaging was done using a 40x water immersion objective (NA = 1.0) and controlled with Zeiss ZEN image acquisition software.

For surveillance experiments, 20-40 frame stacks (1µm step size, dependent on cell size) were acquired at a frequency of one stack per 1 minute for a total duration of 10 minutes. For chemotaxis, 61 frame stacks (1µm step size) were acquired at a frequency of one stack per 2 minutes for a total duration of 60 minutes. For iC3b-bead interaction experiments, 61 frame stacks (1µm step size) were acquired at a frequency of one stack per 2 minutes for a total duration of 120 minutes.

For stimulus application, glass pipettes were fabricated from borosilicate glass capillaries on a horizontal puller (Sutter Instrument, P-80/PC) and fire polished to a final tip diameter of approximately 2µm for chemotaxis experiments and 5µm for bead application. Injections contained Cascade Blue labeled 3000 MW Dextran (ThermoFisher, D7132), aCSF, and 1mM ATP (need Fritz’s ATP here) for chemotaxis experiments or dextran, aCSF, and pHrodo-iC3b opsonized 2μm beads, as previously described, for bead application. For drug treatment, brain slices were incubated in aCSF containing 200µM of CK-666 or the equivalent amount of DMSO for 2 hours prior to being placed into the tissue slice chamber. Drug was not added to the aCSF flowing through the system during imaging but was added to the solution in the glass pipette for injection at the same concentration as incubation. Surveillance videos did not contain a stimulus application.

### Surveillance analysis

Microglia skeletons and morphometrics prepared using automated plugin MotiQ. Original images were first binarized using the mean algorithm, then the close function was used to capture incomplete processes as continuous. MotiQ cropper was applied when appropriate, and thresholder (with no preprocessing or stack processing) and 2D analyzer (no Gauss filter, remove all particles but the largest) were used to generate skeletons and morphology metrics.

### Chemotaxis analysis

Images were measured using Fiji ImageJ to detect length of process travel in maximum projection videos between injection timepoint and reaching the pipette tip. This same process measured how far processes traveled up the inside of the pipette.

### Bead interaction analysis

Beads were counted by hand for color overlap at each the end timepoint. Processes interacting with the beads were measured using ImageJ.

### Statistical Analysis

The Kruskal–Wallis with Dunn multiple comparisons test was used to assess significance in experiments where a normal distribution of the dataset could not be assumed.

When only two experimental conditions were tested, we used Mann–Whitney tests when we could not assume normality. Unpaired t tests and ANOVAs were used when normality tests indicated a normal distribution of the data. All statistics were calculated using GraphPad Prism, and significance was assumed if p ≤ 0.05. More information on each statistical test can be found in the relevant figure legend panel.

---

**Supplementary Figure 1:**
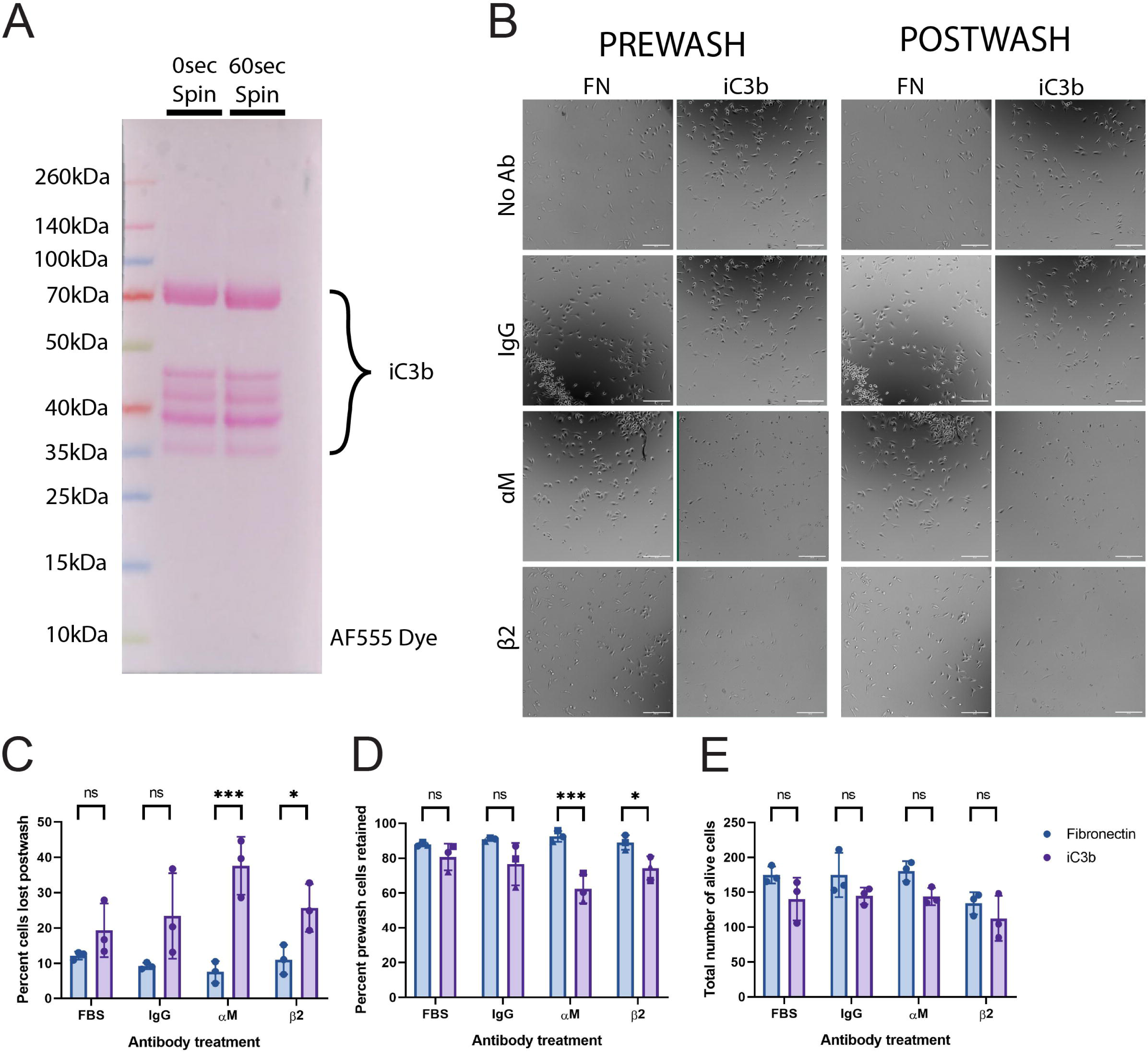
Confirmation of Alexa Fluor 555 labelling of iC3b. A) Ponceau Stain of 8µg of AF-iC3b, either before spinning down or after a 60 second spin down. B) Example images of cells plated on FN or AF-iC3b treated with FBS (vehicle control), IgG (negative control), CD11b, or β2 pre- and post-wash for adhesion assay. Scale bar represents 250µm. C) Percent of cells lost in the process of washing wells in adhesion assay. D) Percent of cells from pre-washing remaining in post-wash images. E) The total number of alive cells in the pre-wash images.

**Supplementary Figure 2:**
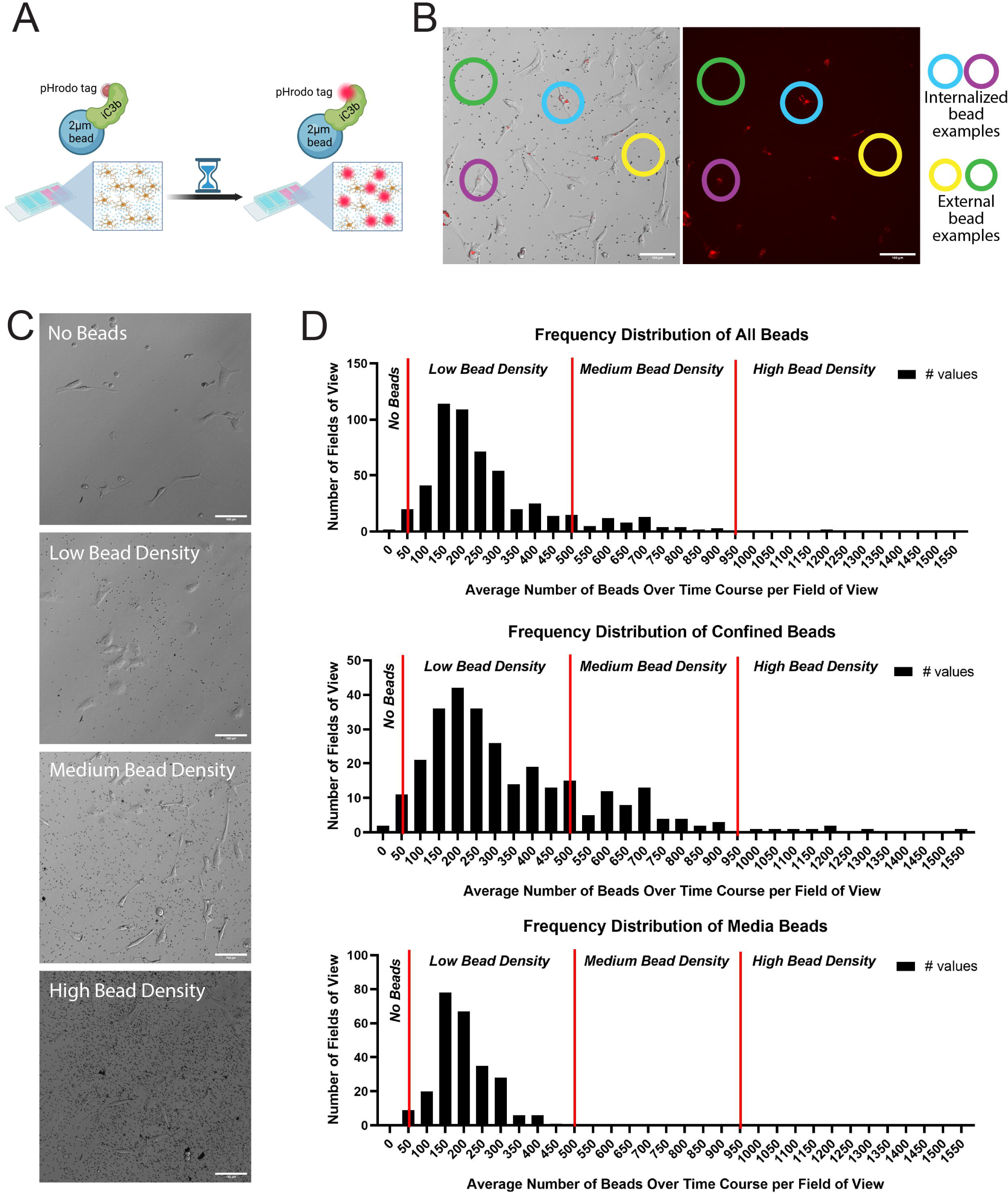
Strategy for controlling for bead density. A) Schematic depicting phagocytic bead labelling and the pHrodo tag fluorescing inside cells but not externally. B) Duplication of Figure 1J. Examples of either internalized (blue and purple circles) or external (yellow and green circles) pHrodo-red staining outlined in both composite and pHrodo only images. Scale bar represents 100µm. C) Example phase contrast images displaying different bead densities. No beads (top) through high bead density (bottom). Scale bar represents 100µm. D) Histograms detailing the breakdown of average bead densities per field of view across all experimental runs. Low bead density was classified as any field of view with a bead average between 50 and 500 beads; medium bead density was 500 to 950 beads; high bead density was any field of view average above 950 beads. Low bead densities in confined images most closely matched the density of typical media images.

**Supplementary Figure 3:**
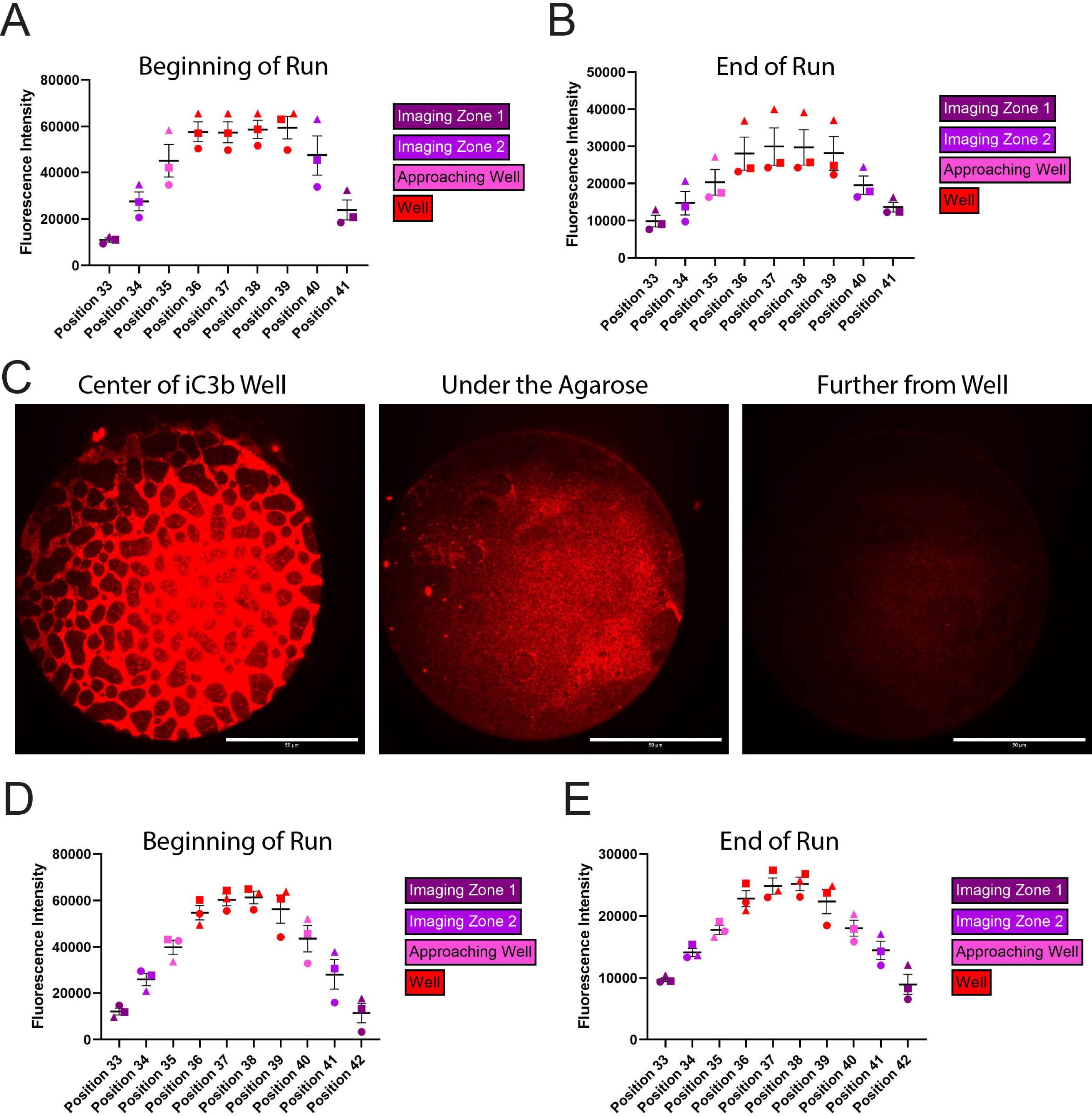
Examining consistency of AF-iC3b labeling during haptotactic assays. A-B) Measurement of the mean fluorescent intensity during haptotaxis runs of the AF-iC3b label spanning from the Zone 1 of one cell well across the center well to the Zone 1 section of the second well. Measurements are color coded by position (see legend) and symbols represent the three haptotaxis runs. (A) corresponds to the beginning of the runs and (B) corresponds to the 24-hour point of the run. C) TIRF images of the glass bottom of the haptotaxis apparatus to demonstrate the binding of AF-iC3b to the glass and the gradient as the signal diminishes the further from the center well the image is taken. Scale bars represent 50µm. D-E) These graphs are the same as (A-B), but with the CK-666 haptotaxis runs.

## ACKNOWLEDGEMENTS

We thank the members of the Rotty Lab for helpful discussions during research development. We thank Dr. Zygmunt Galdzicki for the use of his lab’s vibratome. We thank the Biomedical Instrumentation Center for access to the Zeiss LSM 7MP two-photon microscope. This project is sponsored by the Uniformed Services University of the Health Sciences (USU); however, the information or content and conclusions do not necessarily represent the official position or policy of, nor should any official endorsement be inferred on the part of, USU, the Department of Defense, or the U.S. Government.

## AUTHOR CONTRIBUTIONS

S.P.: experiments and planning, data analysis, writing and editing. I.S..: data analysis, editing. F.L.: experiments and planning, editing. J.R.: Project oversight, experiments and planning, writing, editing, funding. All authors had the opportunity to review and comment on the manuscript prior to submission.

## DATA AVAILABILITY STATEMENT

All primary data will be openly available upon request.

